# Community-based biomedical context to unlock agentic systems

**DOI:** 10.1101/2025.07.21.665729

**Authors:** Malte Kuehl, Darius P. Schaub, Francesco Carli, Lukas Heumos, Camila Fernández-Zapata, Nico Kaiser, Jonathan Schaul, Ulf Panzer, Stefan Bonn, Sebastian Lobentanzer, Julio Saez-Rodriguez, Victor G. Puelles

**Affiliations:** Department of Clinical Medicine, Aarhus University, Aarhus, 8200, Denmark; Department of Pathology, Aarhus University Hospital, Aarhus, 8200, Denmark; Institute of Medical Systems Bioinformatics, Center for Biomedical AI (bAIome), Center for Molecular Neurobiology (ZMNH), University Medical Center Hamburg-Eppendorf, Hamburg, 20251, Germany; III. Department of Medicine, University Medical Center Hamburg-Eppendorf, Hamburg, 20246, Germany; Open Targets, European Bioinformatics Institute (EMBL-EBI), Cambridge, UK; European Molecular Biology Laboratory, European Bioinformatics Institute (EMBL-EBI), Hinxton, UK; Institute of Computational Biology, Computational Health Center, Helmholtz Center, Munich, Germany; Hamburg Center for Kidney Health (HCKH), University Medical Center Hamburg-Eppendorf, Hamburg, 20251, Germany; Institute for Computational Biomedicine, Heidelberg University, Faculty of Medicine, Heidelberg University Hospital, Heidelberg, Germany; Hamburg Center for Translational Immunology (HCTI), University Medical Center Hamburg-Eppendorf, Hamburg, 20251, Germany; German Center for Child and Adolescent Health (DZKJ), partner site Hamburg, University Medical Center Hamburg-Eppendorf, 20251, Germany; Computational Biology Unit, German Center for Diabetes Research, Munich, Germany

**Keywords:** Model Context Protocol, Biomedical AI, BioContextAI, Agentic Systems

## Abstract

Large language models (LLMs) face reliability challenges stemming from hallucinations and insufficient access to validated scientific resources. Existing solutions are often fragmented and limited to specific applications, hindering broader adoption and interoperability. Here, we present Biomedical Context for Artificial Intelligence (BioContextAI), an open-source initiative centered on Model Context Protocol (MCP) servers to address these limitations. BioContextAI provides a community-oriented registry for discovering domain-specific MCP servers and a proof-of-concept server implementation that integrates widely-used biomedical knowledgebases. By enabling standardized access to validated scientific knowledge, BioContextAI aims to facilitate the development of composable agentic systems for biomedical research. Together, this work contributes to an emerging ecosystem of community-driven approaches for expanding the capabilities and reliability of biomedical AI systems.

## Introduction

Large language models (LLMs) have recently been employed in biomedical research for diverse tasks, including cell-type annotation, literature mining, peer review assistance, and knowledge querying [1–3]. Despite their potential applications, LLMs have significant limitations in biomedicine as they often generate hallucinations [4], cannot access specialized databases, and lack the necessary domain knowledge and functional capabilities [5], which are crucial for providing reliable scientific outputs.

Agentic systems have been envisioned as a new paradigm for LLM applications [6] in which agents do not solely rely on model-inherent knowledge but autonomously use available tools to accomplish complex tasks [7]. While web search integration has expanded LLM capabilities, this alone has proved insufficient to realize biomedical research agents, as they require interconnected access to specialized resources such as knowledgebases, medical literature, omics data, and custom applications, often hidden from LLMs behind application programming inter-faces (APIs) not accessed by the crawlers used to generate their training data. This limitation has been recognized [8], and specialized software that leverages retrieval-augmented generation (RAG) [9], custom-developed tool suites [10, 11], or LLMs fine-tuned on biomedical data [12] to integrate previous knowledge has been developed. In addition, artificial intelligence (AI) agents designed to aid with common computational biology analysis flows have begun to emerge [13, 14]. Yet, current solutions are often tailored to specific use cases or only compatible with selected LLM models, applications, or hardware. More-over, a lack of reusable components and open standards places an unnecessary burden on developers of agentic systems to reimplement the layers through which LLMs access domain-specific resources, requiring large-scale efforts and limiting the potential for synthesis and recombination.

In late 2024, the Model Context Protocol (MCP) was introduced [15], aiming to address these challenges by providing a standardized protocol through which generative artificial intelligence (GenAI) systems can use tool calling to interface with external information and applications. A key concept of MCP is the decoupling of tool providers (servers) and LLM-powered tool consumers (clients/hosts), enabling flexible composability (Suppl. Fig. 1) and reducing the effort to build N agentic systems that integrate M functions from N × M to N + M (Suppl. Note 1). This key advantage has been recognized by both the open-source community and commercial LLM providers, and MCP is currently emerging as the *de facto* standard for LLMs to interface with external resources such as APIs [16].

While first biomedical MCP servers have been developed [17, 18], existing implementations are typically only published through GitHub repositories and do not adhere to FAIR4RS [19] principles, an extension of FAIR data [20] principles to research software. MCP partially mitigates interoperability challenges by standardizing the protocol for tool discovery and interaction, but significant gaps remain. The findability of available MCP servers is limited due to the lack of persistent identifiers and a community-based registry that offers rich, searchable metadata. Inconsistent licensing practices, missing metadata, and unclear long-term support further hamper accessibility. Additionally, reusability is impacted by varying documentation quality, unclear data and code provenance, and lack of community development, maintenance, deployment, and integration standards.

## Results

Here, we present **Bio**medical **Context** for **A**rtificial **I**ntelligence (**BioContextAI**), an open-source initiative to index and develop biomedical MCP servers and lay the groundwork for creating composable agentic research assistants (Fig. 1a). At its core, BioContextAI features the **Registry** (Fig. 1b), which adopts an open-source community model inspired by initiatives like the scverse [21], BioCypher [22], Bioconductor [23], and Bioconda [24] (Suppl. Online Methods). It leverages Schema.org ontology terms [25] to catalog community-developed MCP servers in a public repository with rich metadata, enforcing documentation and open-source licensing standards. MCP server developers can use an interactive online editor to seamlessly create metadata files, which can be contributed through the Registry GitHub repository (Suppl. Fig. 2). Researchers can then interact with the Registry through a downloadable index or a searchable interactive web interface available through the BioContextAI web application at https://biocontext.ai/registry.

**Fig. 1.**
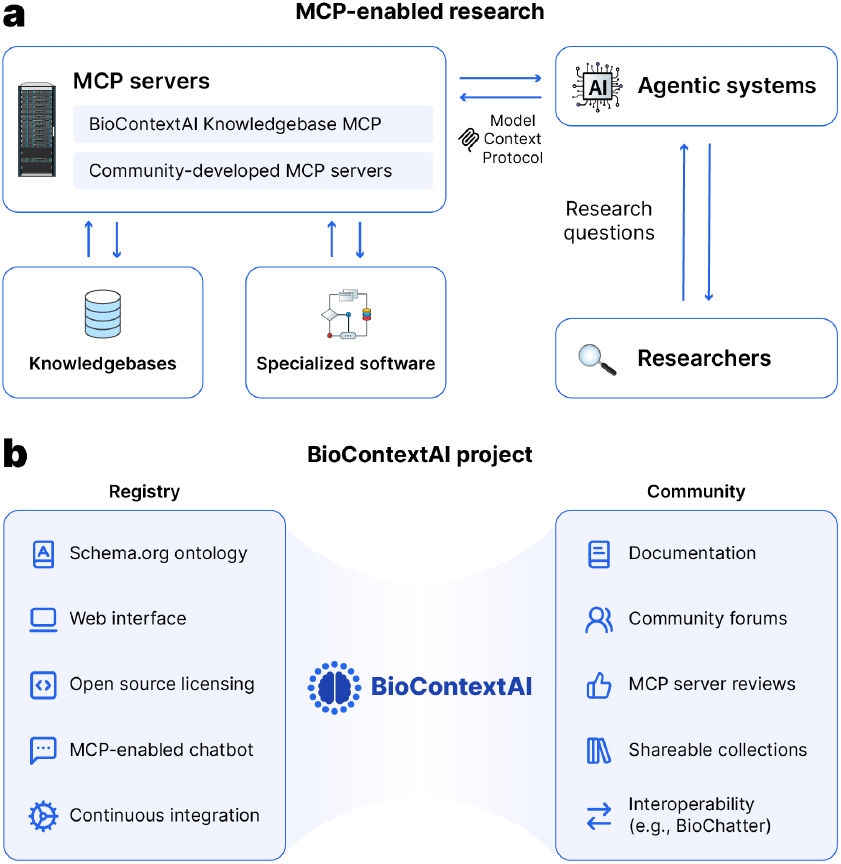
BioContextAI supports community-oriented agentic research in biomedicine. a) Facilitating access to specialized resources. In an Model Context Protocol-enabled paradigm, investigators can pose their research questions in natural language. LLM-based agentic systems can then query MCP servers to gain access to information from knowledgebases or leverage specialized software. **b) Towards a modular MCP server ecosystem**. BioContextAI indexes community-built MCP servers in its Registry with Schema.org ontology, an accessible web interface and related tooling. We further aim to foster a community of MCP server developers and the researchers who use them, offering extensive documentation, forums, a review system and shareable collections to help users choose adequate MCP servers for their scientific endeavors

The web page further enables users to create accounts to freely access **BioContextAI Chat**, an MCP-enabled chatbot integrating different commercial LLMs. In addition, a review system and shareable collections permit the community to identify the most helpful MCP servers for various use cases. Lastly, the documentation at https://biocontext.ai/docs provides useful resources for MCP server development and usage tips for BioContextAI. To provide a foundation for BioContextAI Chat and an example for MCP server development, BioContextAI also offers the **Knowledgebase MCP**, an MCP server implementation that exposes functionality from STRING [26], KEGG [27], PanglaoDb [28], the Human Protein Atlas [29], Reactome [30], Open Targets [31], EuropePMC [32], UniProtKB [33], the Antibody Registry [34], Grants.gov, ClinicalTrials.gov and Drugs@openFDA, resources used for diverse biomedical research applications such as antibody panel design [35] or druggability profiling [36]. It is available as an open-source software package for local deployment and as a remote-hosted server that can be integrated setup-free across an array of GenAI applications. The Knowledgbase MCP server can be installed from the PyPi Python software registry and flexibly combined with existing MCP servers to enhance agentic systems with extended biomedical capabilities.

To demonstrate the potential for MCP servers to improve LLM-assisted biomedical research tasks, we compared the output from Knowledgebase MCP-augmented agentic systems with the respective base and web search-enabled models across different model providers and model sizes (Fig. 2a). This comparison revealed improved response quality with access to the MCP beyond the benefit conferred by web search capability alone (Fig. 2b). We further conducted several case studies (Fig. 2c, Supplementary Data) across a diverse range of topics, including proteinprotein interactions, clinical trials, drug usage information, marker prognosticity and gene-disease associations, providing examples of usefulness for simple and multi-step assignments. To showcase the modularity of MCP-driven agentic system design, we combined the Knowledgebase MCP with a community-developed MCP server to access the OmniPath database [37] within a single LLM conversation (Suppl. Note 2) and provide code to integrate it with BioChatter.

**Fig. 2.**
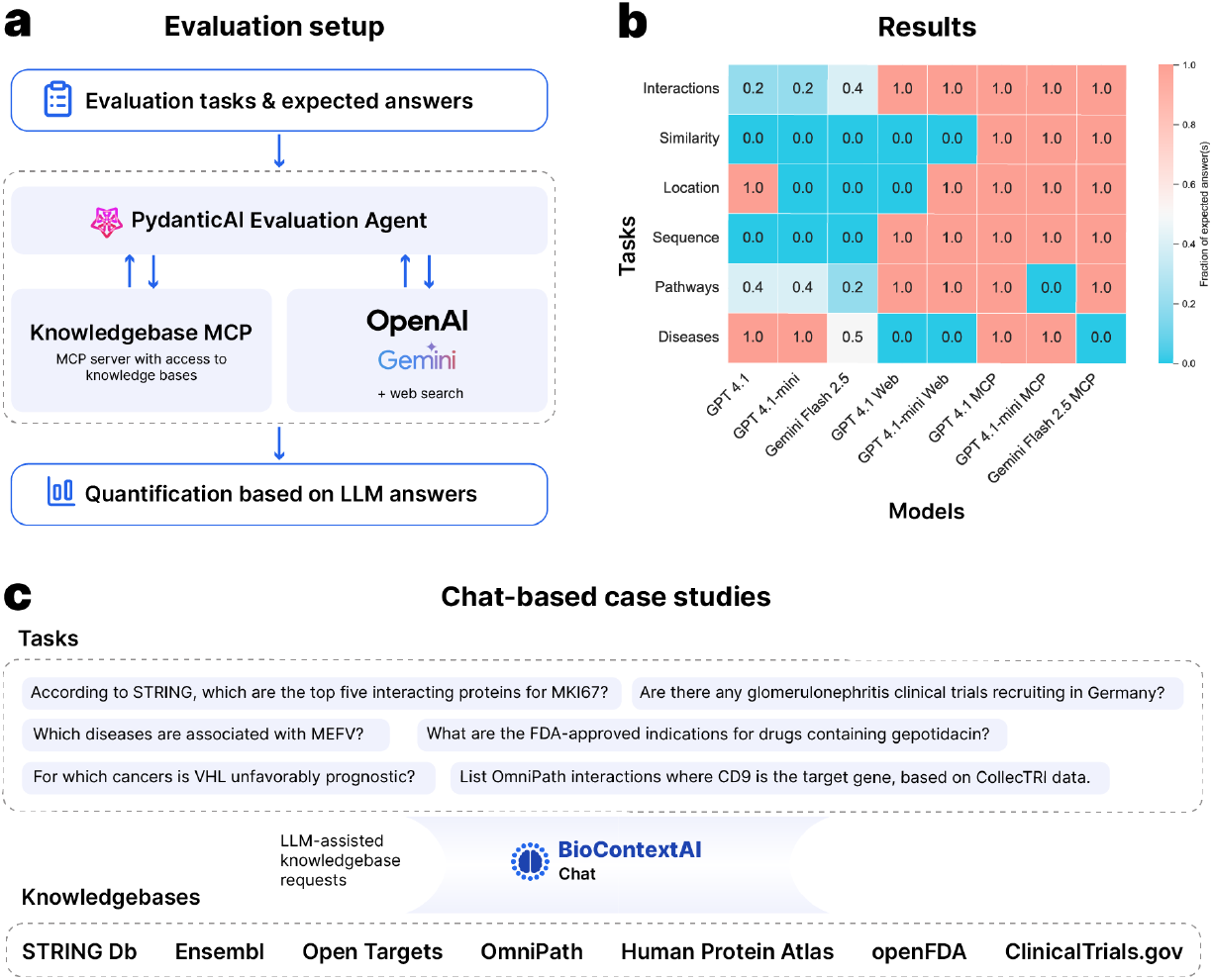
Evaluation and case studies. **a) Setup of the MCP evaluation**. Tasks and expected answers were pre-defined. For each configuration of base model, web search access and MCP server access, a PydanticAI [38] agent was created. **b) Quantification of LLM responses**. LLM answers were compared to the previously defined expected answers across all configurations and tasks, revealing improved responses with MCP server access. Interactions: STRING protein-protein interactions; Similarity: STRING bit similarity score; Location: UniProt subcellular location; Sequence: UniProt amino acid sequence; Pathways: KEGG pathways; Diseases: Open Targets disease associations; GPT: OpenAI GPT; Gemini: Google Gemini; Web: Native web search capabilities; MCP: Access to the BioContextAI Knowledgebase MCP server. **c) Case studies**. To showcase potential use cases for MCP-supported research, we provide additional case studies covering different tasks (Suppl. Note 2). Tasks were evaluated with BioContextAI Chat, resulting in successful access to various knowledgebases exposed via the Knowledgebase MCP and the community-developed OmniPath MCP. The PydanticAI logo is licensed under the MIT license. The OpenAI wordmark and the Google Gemini logo were obtained from Wikipedia under public domain licensing. OpenAI and Gemini remain trademarks of their respective owners.

## Discussion

While the Registry significantly enhances discoverability and constitutes a first step towards establishing FAIR4RS principles, challenges persist. Remote-hosted MCP servers present significant security, authentication, financial, data privacy, and maintenance concerns [39], which puts them out of reach for several research applications. Similarly, state-of-the-art agentic systems often rely on closed-source, closed-weight LLMs, which limits the long-term accessibility and replicability of the academic research that depends on them.

Additionally, evaluating biomedical AI systems is an open research area. While several benchmarks have been proposed [40, 41], actual use cases remain limited, with benchmarks requiring specific data formats [42] or partly focusing on data from a few specialized sources [43]. Furthermore, existing benchmarks primarily evaluate the oveall performance of AI systems. More extensive efforts are needed to profile the impact of their constituent parts in various configurations and their performance on specific sub-domains of scientific research.

Going forward, we are committed to the continued development of BioContextAI to establish the Registry as a community hub for LLM-supported biomedical research, to render further tools available through additional MCP servers, and to expand community resources for creating, evaluating and deploying MCP-augmented agentic systems, through best practices guides, templates, benchmarks and community forums. We invite researchers and developers to engage with BioContextAI by interacting with the community, contributing MCP servers, developing applications, and integrating available tools into research workflows to help advance our understanding of health and disease.

## ACKNOWLEDGEMENTS

We thank Martin Becker for their helpful feedback. This manuscript was partially revised using LLM-based technology. The entire manuscript has been checked for correctness, and the authors are responsible for its final content.

## AUTHOR CONTRIBUTIONS

M.K., D.P.S., and V.G.P. conceived the study. MK wrote the first draft. M.K. and D.P.S. developed the BioContextAI Knowledgebase MCP package, the Registry, BioContextAI Chat and the MCP server evaluation, with F.C., S.L., J.S., C.F.-Z. and N.K. providing inputs. F.C. and S.L. implemented the BioChatter integration. M.K., L.H., D.P.S., S.L., F.C., J.S.-R. and V.G.P. developed the community structure. M.K. and V.G.P. created the figures. U.P., S.B., J.S.-R., S.L., and V.G.P. acquired funding. J.S.-R. and V.G.P. supervised the study. All authors approved and contributed to the editing of the final draft.

## DATA AVAILABILITY

The transcripts of the MCP evaluation queries and responses have been deposited at Zenodo and can be accessed via DOI: https://doi.org/10.5281/zenodo.16163431.

## CODE AVAILABILITY

The BioContextAI Registry is available at: https://github.com/biocontext-ai/ registry. A searchable index and tutorial resources are available at: https://biocontext.ai. The BioContextAI Knowledgebase MCP code is available at: https://github.com/biocontext-ai/knowledgebase-mcp. It can be installed from PyPi as *biocontext-kb*. A remote-hosted version is available with streamable HTTP transport at: https://mcp.biocontext.ai/mcp/ (subject to usage restrictions). The API documentation is available at https://docs.kb.biocontext.ai. BioContextAI Chat is available as a freely usable chat interface at: https://biocontext.ai/chat. Source code for the BioContextAI website – including the Registry interface and Chat – is available at: https://github.com/biocontext-ai/website. The evaluation scripts can be accessed at https://github.com/biocontext-ai/simple-mcp-evaluation. A simple example app to use the Knowledgebase MCP server with BioChatter is provided at: The evaluation scripts can be accessed at https://github.com/biocontext-ai/biochatter-mcp-example.

