## Supplementary Information for "Community-based biomedical context to unlock agentic systems"

### Supplementary Online Methods

BioContextAI is an open-source initiative for integrating biomedical tools and AI systems via the Model Context Protocol (MCP). It includes a community-driven registry of MCP servers for biomedical applications, a modular MCP server that connects to REST- and GraphQL-based knowledgebases, and a hosted chat interface. BioContextAI provides a foundation for flexible tool integration across agentic AI systems and is actively maintained and extensively documented.

**BioContextAI Registry.** The BioContextAI **Registry** is a community-based registry of biomedical MCP servers. The searchable index, based on a GitHub repository, allows tool creators to add their MCP servers via *meta.yaml* files that follow a subset of the Schema.org (version 29.2) SoftwareApplication ontology [1]. The provided schema ensures the presence of relevant metadata, including the tool's name, description, repository URL, maintainers, and open-source license, to help ensure adherence to FAIR4RS principles [2]. The repository can be accessed at https://github.com/biocontext-ai/registry. Developers of MCP servers that may be relevant to biomedical research use are welcome to submit pull requests to have their tools added to the index. They can use the graphical editor provided at https://biocontext.ai/registry/editor to effortlessly create the required metadata file. Upon acceptance of MCP server contributions, an updated version of the Registry list is deployed as a JSON file through continuous integration/continuous deployment (CI/CD). The Registry JSON file is publicly accessible at https://biocontext.ai/registry.json. To provide users with a more intuitive and graphical interface, the registry data is regularly imported into a PostgreSQL database and combined with additional data, including READMEs and star counts obtained via the GitHub API and a list of available tools, their description and input schema, automatically generated from remote-hosted MCP servers. The graphical registry interface is available at https://biocontext.ai/registry. Users also have the opportunity to authenticate through GitHub or LinkedIn via OAuth to mark MCP servers as helpful and create personalized collections of MCP servers. The documentation at https://biocontext.ai/docs contains additional tutorials and links to relevant resources to help potential users and tool developers integrate with the BioContextAI Registry and develop or use novel tools. The source code for the web application is available at https://github.com/biocontext-ai/website.

**BioContextAI Chat.** BioContextAI **Chat** is a Next.js-based agentic chatbot built with the Vercel AI SDK and Model Context Protocol SDK to consume streamable HTTP MCP servers through MCP clients. The application provides text-streaming capabilities and serverless hosting and includes a viewer to review MCP tool calls. Additionally, it features a flexible web interface that allows users to add and configure MCP servers directly within the application. The public deployment of BioContextAI Chat, available at https://biocontext.ai/chat, offers free but rate-limited usage of both Google's Gemini 2.0 Flash-Lite and 2.5 Flash models. Users can optionally utilize OpenAI's GPT-4.1-mini and GPT-4.1 or Anthropic's Claude Sonnet 4 by providing their own API key. Thanks to the comprehensive support of the AI SDK and its abstraction of model providers, developers interested in incorporating different LLM models can easily adapt the source code to meet their specific requirements. The source code is contained within the overall website code and is available at https://github.com/biocontext-ai/website.

**BioContextAI Knowledgebase MCP.** The BioContextAI **Knowledgebase MCP** server facilitates access to several commonly used biomedical knowledgebases and its basic structures acts as an reference implementation for multi-tool community MCP servers. It is based on the Python FastMCP [3] framework. Its structure is highly modular and allows for the flexible integration of three different types of APIs: 1) REST-based APIs with available OpenAPI [4] schemas can be added through minimal configuration via a YAML text file. The Knowledgebase MCP server automatically loads and parses the schema at start-up and exposes all defined endpoints as tools. 2) When REST-based APIs do not follow the OpenAPI specification or high-level functions are desired, tools are implemented through Python functions that use the requests library to access relevant endpoints. 3) Lastly, the MCP server contains code to integrate GraphQL APIs based on their schema URLs. BioContextAI Knowledgebase MCP is unit-tested through pytest, extensively annotated, and tools are typed with Pydantic to provide useful context and constraints to LLMs. It is available as a remote-hosted MCP server through streamable HTTP transport at https://mcp.biocontext.ai/mcp/ (subject to usage and availability restrictions). Furthermore, it can be freely interfaced with through BioContextAI Chat. The source code, including local development and remote deployment instructions, is available at https://github.com/biocontext-ai/knowledgebase-mcp. It can be installed locally from the PyPi Python software repository, where it is available as *biocontext-kb*.

**Evaluation & Case Studies.** To showcase the impact of MCP servers on different research tasks, we created a simple evaluation setup based on PydanticAI [5] designed to interface with LLMs and MCP servers. We established a baseline for each model without MCP access, a web search-enabled version, and an MCP-augmented version that used the BioContextAI Knowledgebase MCP. Our comparison included the following models: OpenAI GPT 4.1-mini (version 2025-04-14), OpenAI GPT 4.1 (version 2025-04-14), and Google Gemini 2.5 Flash (June 2025 update). All models were configured to use a fixed seed of 3407, a maximum token count of 10,000, a temperature of 0.0, and at most 10 steps per message. When tools were provided, models were encouraged to use them through the system prompt. When web search was enabled, models were obligated to use it. Ground truths were manually established based on data available through the APIs and web interfaces of the respective services. Task success was quantified by manually assessing whether the previously established expected answer was provided. Where expected answers had multiple parts, e.g., in the case of gene-pathway associations, a score of 1 divided by the number of expected answers parts was given for each correct element contained in the LLM response. Responses that deviated from the expected answers but were judged to be biologically correct, e.g., providing the mature amino acid sequence instead of the immature one, without specification in the prompt, the response was considered correct. The evaluation was performed on 2025-07-03 using a development preview of version 0.1.0 of the Knowledgebase MCP. In addition to this programmatic execution, we provide further case studies of possible use cases included in Supplementary Note 2. These case studies are intended to explore possible use cases and highlight the interoperability of MCP servers. In addition to the BioContextAI Knowledgebase MCP, they also include an example of a multi-MCP server use case, combining the remote Knowledgebase MCP with a remote Omnipath MCP server (early development version). These case studies were generated through the BioContextAI Chat web interface. Furthermore, we also explored using the Knowledgebase MCP through BioChatter, highlighting how BioContextAI readily integrates with existing specialized solutions for biomedical AI applications. To this end, we linked BioChatter with a Chainlit interface. The evaluation and case study code can be found at `https://github.com/biocontext-ai/simple-mcp-evaluation`. Conversation transcripts for BioContextAI Chat-based interactions are included in Supplementary Note Supplementary Note 2 and evaluation transcripts have been deposited via Zenodo with DOI: `https://doi.org/10.5281/zenodo.16163430`. The BioChatter integration code is provided at: `https://github.com/biocontext-ai/biochatter-mcp-example`.

### Supplementary Figures

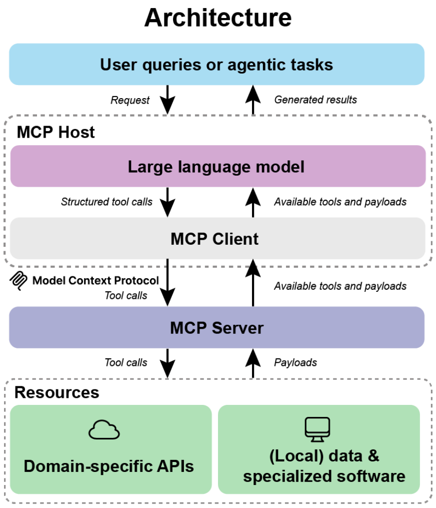


**Fig. S1. Architecture of an MCP-enabled workflow.** In the Model Context Protocol paradigm, queries are processed by artificial intelligence systems that include a large language model and an MCP client. The client connects via MCP to an MCP server (either locally or through network requests) and exposes available functionality to the LLM. Upon tool calls, the structured request is forwarded to the MCP server which – depending on the use case – leverages resources such as external APIs, data access or the ability to execute software. The generated payload is then returned to the LLM to inform its response to the user query. This setup isolates the MCP host (application layer) from the MCP server, allowing for independent development and full reusability of the server, thanks to the standardized protocol. MCP hosts can typically be configured to access multiple MCP servers, allowing for flexible composability of agentic systems.

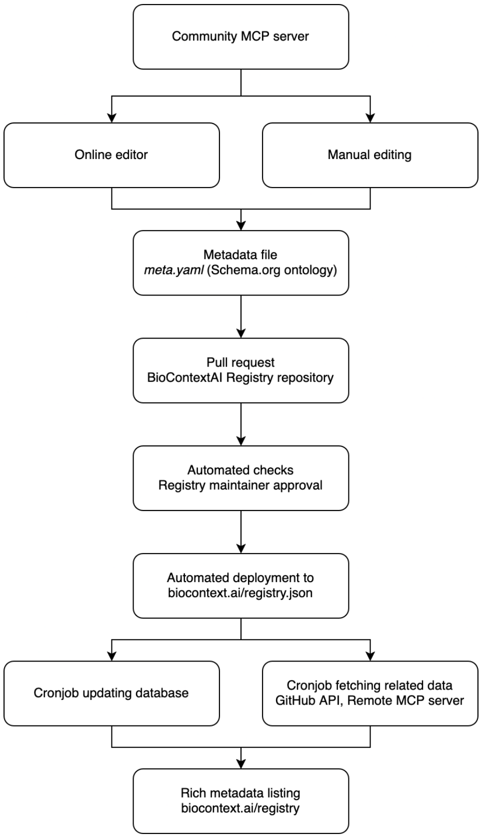


**Fig. S2. Contributing MCP servers to the Registry.** Metadata files for community MCP servers can be generated through the hosted online editor or manually, based on Schema.org ontology terms. These files, along with a checklist for inclusion, are then contributed to the Registry GitHub repository through pull requests. After automated schema validation and approval by Registry maintainers, the MCP servers are added to the publicly accessible Registry JSON database. Cron jobs automatically push these changes to the PostgreSQL database underlying the Registry web interface, combining them with additional data. This includes GitHub repository information obtained through the GitHub API and tool specifications obtained from remote-hosted MCP servers through an MCP client, providing additional information to Registry users.

### Supplementary Note 1: Considerations regarding the Model Context Protocol

**Definitions.** The Model Context Protocol specification defines the role of the MCP host, the MCP client and the MCP server [6]. For the reader's convenience, we summarize the concepts here:

- **MCP Host**: The LLM application (e.g., chatbot application, integrated development environment with AI capabilities, custom agentic system). The host may create and manage one or multiple clients, through which it can connect to MCP servers.
- **MCP Client**: A component within the host application that maintains a 1:1 connection with an MCP server. Clients handle the protocol and relay messages between the host and the server.
- **MCP Server**: Servers provide additional functionality and may expose external data or software. Servers can run locally or remotely, providing different capabilities:
  - Tools: Functions for model-controlled actions, e.g., querying an API.
  - Resources: Context that can be provided to AI models, e.g., documents.
  - Prompts: Interaction templates for users.
  - Sampling: Requests for LLM completions that are sent back to the server.

For further information, we encourage readers to refer to the latest version of the Model Context Protocol specification available at: `https://modelcontextprotocol.io`.

**Motivation for the Use of the Model Context Protocol.** Given that LLMs have supported tool calling for multiple years and that retrieval-augmented generation has been used to provide relevant context to LLMs [7] to accurately answer queries based on factual knowledge and domain-specific information, it may not immediately appear obvious how the Model Context Protocol [8] poses a new paradigm in developing and deploying generative artificial intelligence (GenAI) applications. However, it constitutes a break with the development patterns. Where previously for M models and N tools it would have been necessary to write M × N integrations, the detachment of the tool consumer, i.e., the agentic system (host), from the tool provider (server) reduces this effort to M + N [9]. Similarly, compliance with MCP specifications serves as an integration test, providing increased robustness. Moreover, it fundamentally shifts how generative AI applications can be developed and deployed. Where previously considering the desired capabilities of a system had to be part of the development process of the host, it is now possible to ship applications without having to anticipate their use cases, user base and future tool development, as MCP servers and thus additional functionalities can be linked at run time and independently updated and extended. This shift in how GenAI applications are developed empowers users who are no longer bound to the choices of any one provider but can flexibly compose their applications to fit their needs by connecting hosts to MCP servers.

**Difference Between the Model Context Protocol and Retrieval-Augmented Generation.** The Model Context Protocol is distinct from retrieval-augmented generation (RAG), which is a technique that enhances LLMs by retrieving relevant information from knowledge sources before generating responses [10]. RAG typically involves converting documents or document shards into vector embeddings using encoder models and storing them in specialized databases. Briefly, MCP standardizes the integration of models and tools via a protocol, and RAG provides a methodology for knowledge retrieval. Notably, RAG applications can be implemented using MCP, which enables the incorporation of retrieval mechanisms as tools that LLMs can access dynamically, including over the network. This further demonstrates the interoperability of the MCP pattern, which can leverage tools that leverage concepts such as RAG and integrate them across AI systems without extensive re-architecting efforts.

**Limitations of the Model Context Protocol Paradigm.** The Model Context Protocol is a versatile integration layer that facilitates the composition of GenAI applications and their connection to external APIs and tools. However, it is important to note that there are significant considerations to take into account when constructing MCP servers and MCP-enabled applications. Firstly, the discovery and utilization of tools by LLMs is contingent upon the model's capabilities, the adequacy of tool descriptions, and availability of suitable tools. Secondly, while integrating any external API that adheres to the OpenAPI specification is a relatively straightforward process, such interfaces are often constructed in a highly modular and normalized manner, typically requiring multiple chained requests to carry out common user tasks. This can result in a failure of LLMs to identify the requisite sequence of steps, hindering their ability to execute the intended action. Furthermore, sizable response payloads comprising hundreds or thousands of entries, which could generally be processed promptly on present-day computer hardware using conventional scientific software, might exceed the context length of available LLM models or result in costlier, slower responses. Consequently, the "plug and play" nature of the Model Context Protocol is somewhat constrained to cases where available APIs and tools are already well-suited for use with LLMs or can be converted to possess these properties.

On the topic of token count and cost, we propose that with MCP augmentation, smaller models may potentially be sufficient to accurately answer queries. Our MCP evaluation and case studies, comparing MCP-augmented, web search-enabled and base models across OpenAI GPT-4.1, OpenAI GPT-4.1-mini and Google Gemini 2.5 Flash, provide preliminary evidence supporting this hypothesis (Fig. 2, Supplementary Note 2, Supplementary Data). Although increased token count from extended context represents a significant consideration, this approach could potentially reduce overall compute requirements and associated costs when implemented with smaller and/or specialized tool-calling models, while maintaining comparable factual accuracy. Further research is needed to fully validate these efficiency dynamics across diverse applications and model architectures. Given that complex real-world tasks may necessitate the frequent combination of MCP-enabled information retrieval or tool use and model-inherent reasoning, multi-model systems that dynamically route queries based on tool-calling necessity and task complexity may provide a cost-effective balance between computational efficiency and performance quality.

While hallucinations may be inherent to LLMs [11], combining MCP-augmented LLMs with methods to detect hallucinations [12] could further increase the usefulness of these systems.

Recognizing these challenges and implementing the necessary adaptations could improve the accuracy of MCP-augmented AI systems' responses while making them economically competitive by reducing their computational requirements. These adaptations could also enhance the user experience by swiftly and reliably executing complex research tasks via natural language requests.

### Supplementary Note 2: Case Studies

**Use Cases.** To highlight the utility of employing the BioContextAI Knowledgebase MCP, we provide a number of exemplary use cases. These are intended to illustrate how LLMs can interface with the provided tools to provide factual information from external sources. Real-world applications may include follow-up messages and a higher degree of tool chaining to accomplish a wider range of tasks. We encourage readers to try out MCP servers through BioContextAI Chat at `https://biocontext.ai/chat` to develop an understanding of how these integrations may facilitate their research tasks. As base models evolve and more tools are added to both the BioContextAI Knowledgebase MCP and the broader BioContextAI Registry, additional use cases will become available.

**Supplementary Table 1.** Knowledgebases and example queries

| Knowledgebase | Use case | Example queries |
| --- | --- | --- |
| STRING | Protein-protein interactions | • Which are the top 5 interacting proteins for MKI67, according to STRING?<br>• What is the similarity bit score between ADAM9 and ADAMTS9? |
| PRIDE | Proteomics database | • List recent PRIDE studies looking into CD44. |
| InterPro | Functional protein analysis | • Name predicted domains and sites for human CD9. |
| KEGG [1] | Gene – pathway associations | • Which KEGG pathways include human TRPC6?<br>• Which genes are contained in the cGMP signaling pathway according to KEGG?<br>• Get the drug names for ibuprofen and aspirin, then check for interactions and report which genes are affected, all using KEGG. |
| Reactome | Gene – pathway associations | • Which Reactome pathways include human MTHFR? |
| Ontology Lookup Service | Ontology mapping | • What is the Mondo id for the Tetralogy of Fallot? |
| EuropePMC | Literature research | • According to EuropePMC, which are the most recent articles for Fabry disease?<br>• According to EuropePMC, what has Sarah Teichmann recently published on? |
| PanglaoDb | Marker genes for single-cell RNA-sequencing | • Name 5 sensitive marker genes for Podocytes in humans.<br>• Which cells is SLC5A12 a marker of, according to PanglaoDb? |
| Human Protein Atlas | Protein information | • Summarize the Human Protein Atlas information for BRAF.<br>• Which antibodies are available for human CALB1 according to the Human Protein Atlas?<br>• According to the Human Protein Atlas, for which cancers is VHL unfavorably prognostic? |
| UniProtKB | Functional protein information | • In which subcellular compartment is human PCNX3 expressed according to UniProt?<br>• What is the protein sequence of human CXCL6? |

*Continued on next page*

[1] Available in locally hosted version only due to licensing restrictions

**Supplementary Table 1.** Knowledgebases and example queries (continued)

| Knowledgebase | Use case | Example queries |
|---|---|---|
| Grants.gov | Funding opportunities | • List grant opportunities for kidney research. |
| Open Targets | Target – disease associations | • Use Open Targets to list known drugs for chronic myelogenous leukemia.<br>• Which genes are associated with hyperparathyroidism? |
| Drugs@openFDA | Approved drug information | • What is the maximum daily dosage of ibuprofen?<br>• What are the FDA-approved indications for tocilizumab? |
| ClinicalTrials.gov | Clinical trial details | • Which currently recruiting studies look into sacubitril?<br>• Are there any active clinical trials researching FSGS in Germany? If so, please provide details on the first one. |

**Case Studies.** Here, we show example conversations across models from different LLM providers, comparing performance of base models with their BioContextAI Knowledgebase MCP-augmented counterparts. User messages are shown in blue, tool calls in grey and replies from the assistant in green. The following conversations highlight how MCP augmentation can improve chatbot performance across a range of tasks, including protein-protein interactions (Conversation 1) and prognostic markers (Conversation 2). In use cases where base models already possess relevant knowledge, the access to structured information still adds value to LLM responses, by including identifiers that enable users to access further information more readily and by providing additional details (Conversation 3). Additionally, LLMs do not only act as a way to access existing information but also enable recombination and transformation of data into different presentation formats, such as tables (Conversation 4). When access to relevant source information is provided, MCP augmentation can ensure enhanced veracity and help answer nuanced queries. For example, the drug gepotidacin has been evaluated for use in acute bacterial skin and skin structure infections [13] but has only received FDA approval for the specific indication of uncomplicated urinary tract infections [14] at the time of writing (June 2025). With MCP augmentation, the LLM is able to successfully restrict the provided information to this indication when asked about FDA approved indications, while the base model fails to do so (Conversation 5). Moreover, combining multiple MCP servers, e.g., an MCP server for the OmniPath database [15] and BioContextAI Knowledgebase MCP, enables users to assess complementary information through a unified interface (Conversation 6). Together, these case studies underline the need for the integration of knowledgebases and real time access into GenAI applications to better map user queries to factual information from validated scientific sources.

**Conversation 1.** Knowledge of Protein-Protein Interactions.

**Google Gemini 2.0 Flash Lite – MCP server(s): BioContextAI Knowledgebase MCP**

According to STRING, which are the top five interacting proteins for MKI67?

Call to *bc_get_string_interactions*

The top five interacting proteins for MKI67 according to STRING, with a minimum combined score of 900, are:

1. NIFK (score: 0.996)
2. TOP2A (score: 0.993)
3. CENPF (score: 0.979)
4. PPP1CC (score: 0.964)
5. BUB1 (score: 0.963)

5/5 correct.

**Google Gemini 2.0 Flash Lite – MCP server(s): None**

According to STRING, which are the top five interacting proteins for MKI67?

I can help you with that. Here are the top five interacting proteins for MKI67 according to STRING:
**Top 5 Interacting Proteins for MKI67**
Based on the STRING database (`https://string-db.org/`), the top five interacting proteins for MKI67 (also known as Ki-67) are:

1. PCNA (Proliferating Cell Nuclear Antigen)
2. CBX3 (Chromobox Protein Homolog 3)
3. HMGB2 (High Mobility Group Box 2)
4. NCL (Nucleolin)
5. CDK1 (Cyclin Dependent Kinase 1)

**Note:** The STRING database provides a comprehensive resource for known and predicted protein-protein interactions. The interactions are based on various sources, including experimental data, co-expression, and text mining.

0/5 correct

**Actual**[2]
NIFK (0.996)
TOP2A (0.993)
CENPF (0.979)
PPP1CC (0.964)
BUB1 (0.963)

[2]Source: `https://version-12-0.string-db.org/cgi/network?networkId=bIYVrTaCanLO` (Date of consultation: 2025-06-19)

**Conversation 2.** Knowledge of Prognostic Markers.

**OpenAI GPT 4.1 – MCP server(s): BioContextAI Knowledgebase MCP**

According to the Human Protein Atlas, for which cancers is VHL unfavorably prognostic?

Call to *bc_get_human_protein_atlas_info*

**VHL Unfavorable Prognostic Cancers (Human Protein Atlas)**
According to the Human Protein Atlas, VHL (Von Hippel-Lindau tumor suppressor) is unfavorably prognostic for:

- **Liver Hepatocellular Carcinoma (TCGA)**
  - Prognostic type: Unfavorable
  - Statistical significance: p = $6.76 \times 10^{-9}$

This means higher or altered expression of VHL is associated with a worse prognosis in liver hepatocellular carcinoma, based on TCGA data.
**Note:** For other cancers in the Human Protein Atlas, VHL is either not prognostic or is not associated with unfavorable prognosis.

---

Source: Human Protein Atlas (https://www.proteinatlas.org/) If you need details for other cancer types or validation cohorts, let me know.

1/1 correct.

**OpenAI GPT 4.1 – MCP server(s): None**

According to the Human Protein Atlas, for which cancers is VHL unfavorably prognostic?

**VHL (Von Hippel-Lindau tumor suppressor) – Prognostic Summary from the Human Protein Atlas**

**Unfavorable Prognostic Cancers**
According to the Human Protein Atlas (HPA), **VHL** is considered an **unfavorable prognostic marker** for the following cancers:

- **Renal cancer** (kidney cancer)

**Details:**

- **Unfavorable prognostic marker** means that higher expression of VHL is associated with **worse overall survival** in patients with renal cancer, based on HPA RNA expression data and survival analysis.

**References:**

- VHL – Pathology summary, Human Protein Atlas

0/1 correct.

**Actual**[3]
Liver cancer (LIHC)

[3] Source: https://www.proteinatlas.org/ENSG00000134086-VHL/cancer (Date of consultation: 2025-06-19)

**Conversation 3.** Knowledge of Disease Associations.

**OpenAI GPT 4.1 – MCP server(s): BioContextAI Knowledgebase MCP**

Which diseases are associated with MEFV?

Call to *bc_get_ensembl_id_from_gene_symbol*

Call to *bc_get_open_targets_query_examples*

Call to *bc_query_open_targets_graphql*

**Diseases Associated with MEFV**

The MEFV gene (Ensembl ID: ENSG00000103313) is associated with a wide range of diseases, most notably autoinflammatory and genetic disorders. Below are some of the most strongly associated diseases, based on Open Targets data and their respective ontology IDs:

| Disease Name | Ontology ID | Evidence Score (max) |
|---|---|---|
| Familial Mediterranean fever | MONDO_0018088 | 0.94 (EVA) |
| Autosomal recessive familial Mediterranean fever | MONDO_0009572 | 0.99 (UniProt) |
| Sweet syndrome | MONDO_0011959 | 0.92 (EVA) |
| Autoinflammatory syndrome | MONDO_0019751 | 0.89 (EVA) |
| Behcet's syndrome | EFO_0003780 | 0.74 (EVA) |
| Ankylosing spondylitis | EFO_0003898 | 0.57 (GWAS) |
| Congenital heart disease | EFO_0005207 | 0.44 (EVA) |
| Neuronal ceroid lipofuscinosis | MONDO_0016295 | 0.44 (EVA) |
| Lung adenocarcinoma | EFO_0000571 | 0.69 (Slapenrich) |
| Ovarian cancer | MONDO_0008170 | 0.61 (Slapenrich) |
| Systemic lupus erythematosus | MONDO_0007915 | 0.19 (EVA) |
| Urticaria | EFO_0005531 | 0.18 (EVA) |

**Other Notable Associations**

- Genetic disorder (EFO_0000508)
- Abnormal heart morphology (HP_0001627)
- Renal insufficiency (HP_0000083)
- Neurodevelopmental delay (HP_0012758)
- Hyperpigmentation of the skin (MONDO_0019289)
- Skin vesicle (HP_0200037)
- Syncope (HP_0001279)
- Red cell distribution width (EFO_0009188)
- Mean corpuscular hemoglobin (EFO_0004527)
- Lymphocyte count (EFO_0004587)

**Notes**

- The evidence scores reflect the strength of association from various databases (EVA, UniProt, GWAS, etc.).
- Familial Mediterranean fever is the most strongly and consistently associated disease with MEFV.
- MEFV is also implicated in a spectrum of autoinflammatory and immune-related conditions.

**Data Source**

- **Data Source:** Open Targets Platform (https://platform.opentargets.org/)

If you need more detailed information on a specific disease association or the underlying evidence, please specify!

Date of consultation: 2025-06-19.

**OpenAI GPT 4.1 – MCP server(s): None**

Which diseases are associated with MEFV?

**MEFV Gene: Disease Associations**
The **MEFV** gene encodes pyrin, a protein involved in the regulation of inflammation. Mutations in MEFV are primarily associated with autoinflammatory disorders.

**Established Disease Associations**
**1. Familial Mediterranean Fever (FMF)**

- **Description**: An autosomal recessive autoinflammatory disease characterized by recurrent episodes of fever and serosal inflammation (peritonitis, pleuritis, arthritis).
- **Prevalence**: Most common in people of Mediterranean descent (e.g., Turkish, Armenian, Arab, Jewish populations).
- **Pathophysiology**: Mutations in MEFV lead to dysregulation of the inflammasome and excessive production of interleukin-1β (IL-1β).

**2. Other Reported Associations**
While FMF is the primary disease, MEFV variants have been reported (with less robust evidence) in association with:

- **Systemic juvenile idiopathic arthritis (sJIA)**
- **Behçet's disease**
- **Rheumatoid arthritis** (as a modifier)
- **Other periodic fever syndromes** (rarely, as a modifier or in atypical cases)

*Note*: For these conditions, MEFV mutations may act as susceptibility factors or modifiers rather than direct causes.

**References & Databases**

- **OMIM:** MEFV (OMIM #608107)
- **GeneCards:** MEFV Gene
- **Orphanet:** Familial Mediterranean Fever (ORPHA:342)

**Summary:**
The MEFV gene is most strongly associated with Familial Mediterranean Fever (FMF), and may play a role as a modifier in other autoinflammatory and autoimmune diseases.

Date of consultation: 2025-06-20.

**Conversation 4.** Knowledge of Clinical Trials.

**Google Gemini 2.5 Flash – MCP server(s): BioContextAI Knowledgebase MCP**

Are there any glomerulonephritis clinical trials recruiting in Germany? Format as a table, grouped by study.

Call to *bc_get_recruiting_studies_by_location*

Here are the glomerulonephritis clinical trials currently recruiting in Germany:

| NCT ID | Brief Title | Overall Status | Conditions | Lead Sponsor | Recruiting Locations in Germany |
|---|---|---|---|---|---|
| NCT06047171 | Rescue of Nephrons With ALE.F02 (RENAL-F02) | RECRUITING | Glomerulonephritis Rapidly Progressive | Alentis Therapeutics AG | Berlin, Dresden, Essen, Hanover, Koeln, Leipzig, Luebeck, München |
| NCT06564142 | Evaluation of Efficacy of Povetacicept in Adults With Immunoglobulin A Nephropathy (IgAN) | RECRUITING | Immunoglobulin A Nephropathy | Alpine Immune Sciences Inc, A Subsidiary of Vertex | Aachen, Berlin, Dresden |
| NCT05268289 | Study of Efficacy and Safety of LNP023 in Participants With Active Lupus Nephritis Class III-IV, +/- V | RECRUITING | Lupus Nephritis | Novartis Pharmaceuticals | Braunschweig, Koeln, Berlin, Ludwigshafen, Mainz, Muenchen |
| NCT04557462 | A Rollover Extension Program (REP) to Evaluate the Long-term Safety and Tolerability of Open Label Iptacopan/LNP023 in Participants With Primary IgA Nephropathy | RECRUITING | Primary IgA Nephropathy | Novartis Pharmaceuticals | Essen, Mainz, Ulm |
| NCT06581198 | A Study of Rapcabtagene Autoleucel in Systemic Lupus Erythematosus (SLE) Patients With Active, Refractory Lupus Nephritis (LN) | RECRUITING | Lupus Erythematosus, Systemic; Lupus Nephritis | Novartis Pharmaceuticals | Leipzig, Essen, Jena, Mainz, Ulm |
| NCT05138133 | Phase 3 Study of Anifrolumab in Adult Patients With Active Proliferative Lupus Nephritis | RECRUITING | Lupus Nephritis | AstraZeneca | Dresden, Düsseldorf, Hannover, Heidelberg, Kirchheim, Köln, Mainz Am Rhein, München, Villingen-Schwenningen |
| NCT05797610 | A Study to Evaluate the Efficacy and Safety of Sefaxersen (RO7434656) in Participants With Primary Immunoglobulin A (IgA) Nephropathy at High Risk of Progression | RECRUITING | Primary IgA Nephropathy | Hoffmann-La Roche | Aachen, Berlin, Erlangen, Essen, Greifswald, Hannover, Köln, Mainz, München, Villingen-Schwenningen |
| NCT03955445 | Long-term Efficacy, Safety and Tolerability of Iptacopan in C3G or IC-MPGN | RECRUITING | C3 Glomerulopathy; Immune-complex-membranoproliferative Glomerulonephritis | Novartis Pharmaceuticals | Essen, Hamburg, Mainz |
| NCT06291376 | Study of Ravulizumab in Immunoglobulin A Nephropathy (IgAN) | RECRUITING | Immunoglobulin A Nephropathy; IgAN | Alexion Pharmaceuticals, Inc. | Aachen, Berlin, Erlangen, Essen, Greifswald, Hannover, Köln, Mainz, München, Villingen-Schwenningen |
| NCT05126277 | Safety, Efficacy and Tolerability of Ianalumab Versus Placebo, Combination With SoC Therapy, in Participants With Active Lupus Nephritis | RECRUITING | Lupus Nephritis | Novartis Pharmaceuticals | Berlin, Dresden, Essen, Mainz |
| NCT06925542 | A Safety and Efficacy Study Evaluating CTX112 in Adult Subjects With Refractory Autoimmune Disease | RECRUITING | SLE (Systemic Lupus); Lupus Erythematosus, Systemic; Lupus Nephritis; Systemic Sclerosis; Inflammatory Myopathy, Idiopathic; Myositis; Diffuse Cutaneous Systemic Sclerosis | CRISPR Therapeutics | Augsburg, Hannover |
| NCT05755386 | Study of Efficacy and Safety of Iptacopan in Participants With IC-MPGN | RECRUITING | IC-MPGN | Novartis Pharmaceuticals | Berlin, Dresden, Essen, Mainz |
| NCT06342960 | KYSA-3: A Study of Anti-CD19 Chimeric Antigen Receptor T-Cell (CD19 CAR T) Therapy, in Subjects With Refractory Lupus Nephritis | RECRUITING | Lupus Nephritis; Lupus Nephritis - WHO Class III; Lupus Nephritis - WHO Class IV | Kyverna Therapeutics | Berlin, Dresden, Düsseldorf, Erlangen, Frankfurt, Hamburg |
| NCT03949972 | The FOrMe Registry (The German Focal Segmental Glomerulosclerosis and Minimal Change Disease Registry) | RECRUITING | Glomerulosclerosis, Focal Segmental; Minimal Change Disease; Idiopathic Nephrotic Syndrome | University of Cologne | Cologne, Berlin, Bonn, Dachau, Essen, Heidelberg, Leipzig, Marburg, Münster |

Date of consultation: 2025-06-20. Without the MCP server, no studies are returned.

**Conversation 5.** Knowledge of Approved Drugs.

**OpenAI GPT 4.1-mini – MCP server(s): BioContextAI Knowledgebase MCP**

According to the label information, what are the FDA-approved indications for drugs containing gepotidacin? What is the typical dosage?

Call to *bc_search_drugs_fda*

Call to *bc_get_drug_label_info*

According to the FDA label information for drugs containing gepotidacin (brand name BLUJEPA):
**FDA-Approved Indications:**

- BLUJEPA is indicated for the treatment of female adult and pediatric patients 12 years of age and older weighing at least 40 kg with uncomplicated urinary tract infections (uUTI) caused by susceptible microorganisms including Escherichia coli, Klebsiella pneumoniae, Citrobacter freundii complex, Staphylococcus saprophyticus, and Enterococcus faecalis.

**Typical Dosage:**

- The recommended dosage of BLUJEPA is 1,500 mg (two 750 mg tablets) taken orally twice daily (approximately 12 hours apart) for 5 days.
- BLUJEPA tablets should be taken after a meal to reduce the possibility of gastrointestinal intolerance.

If a dose is missed, it should be taken as soon as possible, but do not double the dose to make up for a missed dose. For more detailed information, you can refer to the official FDA label: BLUJEPA Label PDF.
Let me know if you need additional details from the label.

**OpenAI GPT 4.1-mini – MCP server(s): None**

According to the label information, what are the FDA-approved indications for drugs containing gepotidacin? What is the typical dosage?

**FDA-Approved Indications for Drugs Containing Gepotidacin:**

As of the latest label information, drugs containing gepotidacin are FDA-approved for the treatment of:

- **Uncomplicated urinary tract infections (uUTI)** caused by susceptible strains of designated microorganisms.
- **Acute bacterial skin and skin structure infections (ABSSSI)** caused by susceptible bacteria.

**Typical Dosage**

- For **uncomplicated urinary tract infections (uUTI)**:
The typical dosage is **1500 mg orally twice daily for 5 days**.

- For **acute bacterial skin and skin structure infections (ABSSSI)**:
The typical dosage is **1500 mg orally twice daily for 7 to 10 days**.

**Notes**

- Dosage may vary based on clinical judgment, patient factors, and specific product formulations.
- Always refer to the specific drug label for detailed prescribing information, including contraindications and warnings.

Date of consultation: 2025-06-20.

**Conversation 6.** Knowledge of Interactions (Multi-MCP).

**OpenAI GPT 4.1-mini – MCP server(s): BioContextAI Knowledgebase MCP, OmniPath MCP**

List OmniPath interactions where CD9 is the target gene, based on CollecTRI data.

Call to *execute_sql_query_on_omnipath_db*

There are 23 OmniPath interactions where CD9 is the target gene based on CollecTRI data. Here are some key details of these interactions:

- The sources are mostly protein complexes involving transcription factors such as FOSL1_JUNB, FOS_JUN, FOS_JUNB, FOSL2_JUND, FOSL2_JUN, JUN, FOSB_JUNB, JUNB, FOSL2_JUNB, JUND, JUN_JUND, JUNB_JUND, FOSL1_JUND, FOSL1_JUN, FOSB_JUND, FOSB_JUN, and PITX2.
- The interactions are transcriptional and mostly represent activation (default activation or regulon).
- The references for these interactions include PubMed IDs such as 16000878, 19381435, 16113057, 8760289, and 11950938.
- Some interactions have additional evidence from DoRothEA, HTRIdb, NTNU.Curated, ExTRI, GEREDB, and ReMap databases.
- The interactions are directed and stimulatory, with no inhibition reported.
- The source and target species are human (NCBI taxonomy ID 9606).

If you want, I can provide a detailed list or specific information about any of these interactions.

Using STRING, identify the protein-protein interactions for CD9.

Call to *bc_get_string_interactions*

The protein-protein interactions for CD9 in human (species 9606) according to STRING with high confidence scores include:

1. CD9 - PTGFRN (score: 0.999)
2. CD9 - IGSF8 (score: 0.999)
3. CD9 - CD81 (score: 0.997)
4. CD9 - TSG101 (score: 0.995)
5. CD9 - CD63 (score: 0.994)
6. CD9 - ITGB1 (score: 0.989)
7. CD9 - CD44 (score: 0.985)
8. CD9 - CD82 (score: 0.983)
9. CD9 - HBEGF (score: 0.980)
10. CD9 - ITGA3 (score: 0.969)

These interactions have various supporting evidence scores indicating strong confidence in their association. If you need more details or additional interactions, please let me know.

Date of consultation: 2025-06-20.
